# A Modular Synthetic Hydrogel with Cell-scale Micropores Modulates Osteocyte-like Morphogenesis and Osteogenic Differentiation *In Vitro*

**DOI:** 10.64898/2026.08.04.742693

**Authors:** Marion Horrer, Doris Zauchner, Mélanie Escudero, Eirini Klinaki, Pei Jin Lim, Marianne Rohrbach, Cecilia Giunta, Ralph Müller, Xiao-Hua Qin

## Abstract

One crucial step during early osteogenesis is the embedding of osteoblasts within a collagen-rich extracellular matrix (osteoid), where they subsequently differentiate into a functional network of osteocytes. However, reconstructing 3D osteocyte networks *in vitro* remains a major challenge. We recently developed a synthetic microporous hydrogel to support the *in vitro* culture of 3D bone cell networks. Although matrix biodegradability facilitates cell-material interactions, the influence of micropores on bone tissue morphogenesis and differentiation remains poorly understood. Here, we investigate the effect of cell-scale micropores on bone cell morphogenesis and osteogenic differentiation *in vitro*. By exploiting polymerization-induced phase separation (PIPS) between 4-arm polyethylene glycol vinyl sulfones and dextran in the presence of hyaluronan, we generated matrix metalloproteinase-sensitive hydrogels with cell-scale micropores. Increasing the dextran concentration enlarged the average pore size from 4 μm to 8 μm, accompanied by a slight decrease in mechanical stiffness. Following encapsulation within these hydrogels, primary human osteoblasts remained highly viable. Hydrogels with larger pores supported extensive 3D cell network formation, whereas hydrogels with smaller pores exhibited enhanced osteogenic differentiation following 21 days of osteogenic culture. Together, these findings highlight that bone cells are sensitive to microporous physical cues and even minor changes over pore sizes can make an impact on osteocyte-like morphogenesis and differentiation *in vitro*.

## 1. Introduction

The early stage of osteogenesis is a dynamic process in which bone-forming osteoblasts deposit a collagen-rich osteoid tissue and then a subset of these cells differentiate into a functional network of osteocytes.^1,2^ Recent advances emphasize that various physical cues of the extracellular matrix (ECM), including matrix stiffness, and viscosity determine cell fate and behavior by directing proliferation, differentiation, migration, and apoptosis during early osteogenesis.^3–6^ Skeletal disorders such as osteogenesis imperfecta are characterized by altered matrix development, resulting in hypermineralization and brittleness.^7–9^ Understanding how different ECM cues influence bone development is key for creating accurate *in vitro* models of healthy and diseased bone tissues.

Increasing efforts have been devoted to generating *in vitro* three-dimensional (3D) osteocyte cultures using a variety of hydrogels, including collagen^10^ and fibrin^11^, gelatin methacryloyl (GelMA),^12^ and clickable polyethylene glycol (PEG) or polyvinyl alcohol^13–16^. For instance, Garrigle et al.^17^ studied the impact of matrix stiffness on osteocytic differentiation of MC3T3-E1 murine osteoblasts embedded within gelatin-based hydrogels *in vitro*. The authors demonstrate that softer hydrogels (∼0.6 kPa) are more permissive for osteocyte differentiation and the formation of a cellular network following 56 days.^17^ Several other studies further suggest that lower hydrogel stiffness facilitates osteogenic differentiation of human mesenchymal stromal cells (hMSCs) in bioprinted hydrogels made of alginate-gelatin or GelMA.^15,18,19^

Since native tissues such as osteoid are not purely elastic but viscoelastic, Bernero et al. studied the impact of matrix viscoelasticity on osteogenic differentiation of IDG-SW3 murine osteocytes using an interpenetrating alginate-collagen hydrogel.^20^ While faster relaxing IPN hydrogels foster early osteocyte morphogenesis, slower relaxing hydrogels enhanced osteogenic differentiation following a culture of 14 days.^20^ Beyond matrix mechanics, pore size and porosity have long been recognized as a critical parameter

in bone tissue engineering. Studies using macroporous collagen-GAG scaffolds have demonstrated that pore sizes in the range of 100-150 μm strongly influences cell adhesion, proliferation, and osteogenic differentiation.^21,22^ Notably, these findings were obtained by seeding MC3T3-E1 cells atop the rigid scaffolds.^21,22^ However, how pore sizes influence cell behaviors after they are encapsulated inside a soft osteoid-like matrix remains largely unexplored. Recently, Zauchner et al.^13^ reported a synthetic PEG microporous hydrogel by polymerization-induced phase separation (PIPS) for *in vitro* culture of 3D bone cell networks from hMSCs and human osteoblasts (hOBs). While adding sufficient dextran to the PIPS hydrogel is key to successful cellular network formation, how fine-tuned cell-scale microporosity modulates osteoblast-to-osteocyte transition and osteocyte morphogenesis remains to investigated.

The objective of this study is thus to investigate how cell-scale microporosity in the synthetic matrix metalloproteinase (MMP)-sensitive hydrogel influences the behaviors of 3D embedded hOBs during *in vitro* osteogenic culture. These microporous hydrogels were formed through PIPS between 4-arm PEG vinylsulfone (4-PEG-VS) and dextran using hyaluronic acid (HA) as viscosity enhancer and a MMP-sensitive di-cysteine peptide as crosslinker (**Figure 1**).^13,23^ By varying the dextran concentration, we generated hydrogels with distinct cell-scale microporosities, while maintaining matrix biodegradability and adhesiveness constant.^13,23^ We screened the impact of enzymatic digestion of HA on the microporosity of PIPS hydrogels. By following osteogenic differentiation over 21 days of culture, our findings highlight that even minor adjustment of the micropores can influence osteocyte-like morphogenesis and differentiation within defined hydrogels *in vitro*.

**Figure 1.**
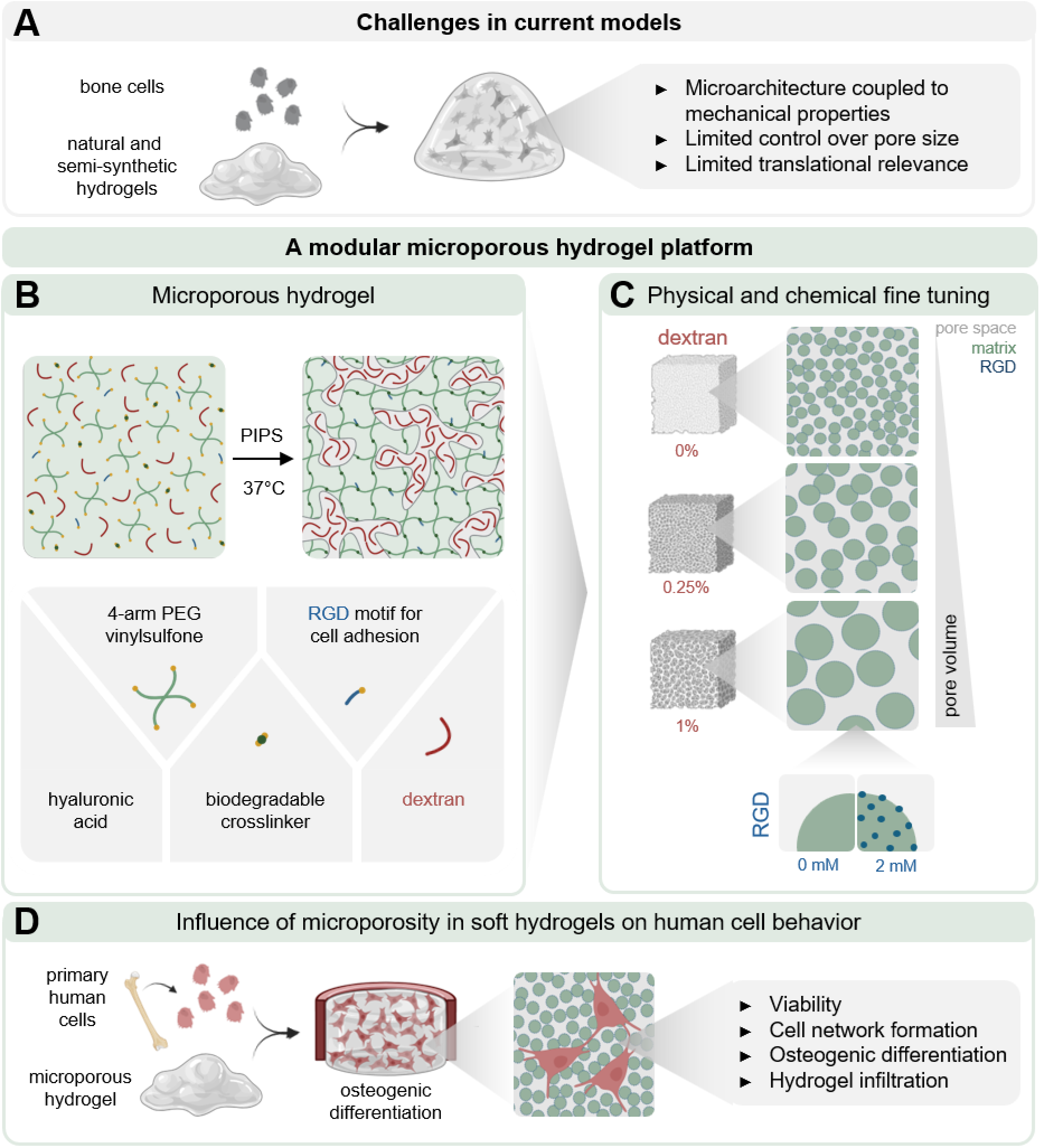
*In vitro* culture of human osteoblasts in modular synthetic microporous hydrogels. **(A)** Schematics of challenges in current models. **(B)** Schematic illustration of the components of porous PEG hydrogels and the concept of polymerization induced phase separation (PIPS). **(C)** Schematic illustration of the influence of varying the dextran concentration on matrix architecture. **(D)** Schematic illustration of the workflow in this study and readouts. Illustration created with Biorender.com (https://biorender.com/djrsq27).

## 2. Results and Discussion

### 2.1. Effect of polymer composition on pore size, swelling and mechanical properties of PIPS hydrogels

In this study, we investigated how fine-tuned pore geometry of the hydrogel matrix influences bone cell fate by varying the composition of a biodegradable microporous hydrogel. After mixing the components at 37°C, the Michael addition crosslinking leads to phase separation, rendering the initially miscible mixture incompatible. This dynamic process uniquely enables one to generate micropores within a hydrogel in the presence of living cells.^13,23^ By varying the dextran concentration within the composition, the PIPS and resulting pore geometry can be modulated accordingly (**Figure 1**).

One key requirement in applying PIPS hydrogels for 3D cell culture is to determine the exact hydrogel compositions that remain miscible (one phase) before crosslinking and subsequently form cell-scale micropores (two phases) *in situ* via PIPS. We thus screened hydrogel formulations for PIPS by varying dextran concentrations from 0% to 4%. For 0-3% dextran, the formulations remain miscible. However, when the concentration of dextran exceeds 3%, the formulations became immiscible, resulting in macroscopically visible phase separation (**Supplementary Figure 1A**). To visualize the hydrogel microarchitecture, we incorporated rhodamine-labeled 4-PEG-VS into the precursor solution. Confocal imaging data confirm that hydrogels with 0.5%-3% dextran showed homogeneous micropores, whereas the hydrogel with 4% dextran exhibit non-homogeneous micropores (**Supplementary Figure 1A-B**). Increasing dextran concentration increased pore size, although only marginal increases were observed above 0.5% dextran (**Supplementary Figure 1B**). Based on these findings, we selected hydrogel formulations containing 0%, 0.25%, and 1% dextran for 3D osteogenic culture of hOBs.

To characterize the pore architecture of the selected formulations, we perfused the equilibrated hydrogels for 4 days with FITC-dextran tracers (*M*_*w*_: 2 MDa) (**Figure 2A**). Confocal imaging revealed uniform tracer distribution across all three groups (**Figure 2B**), confirming interconnected microporous network. Additionally, hydrogels prepared from five independent dextran stock solutions showed consistent micropore sizes (**Supplementary Figure 1C**), indicating the reliability of this PIPS hydrogel platform for reproducible 3D cell culture. Quantitative pore analysis of the FITC-dextran stained hydrogels was conducted using two complementary approaches: 3D sphere fitting analysis^24^ and 2D Fast Fourier transform analysis (RIFFT)^25^. Both methods demonstrated significant increases in pore diameter with increasing dextran concentration (**Figure 2C-H**) and showed strong correlation (*R*^2^ *0*.*99*) (**Supplementary Figure 2A**). 3D analysis results revealed average diameters of 2.8 μm, 4.3 μm, and 5 μm for hydrogels containing 0%, 0.25%, and 1% dextran, respectively (**Figure 2C-D**). In contrast, 2D-RIFFT analysis yielded pore sizes approximately 2-fold larger than those obtained from the 3D analysis (**Figure 2E**). **Figure 2F** shows a slight decrease of porosity from 46% to 44% and 42% as the dextran concentration increases from 0% to 0.25% and 1%, respectively. Pore connectivity analysis confirms an interconnected pore network, with smaller pores predominating at 0% dextran and larger pores at higher dextran concentrations (**Figure 2G-H, Supplementary Figure 2B**).

**Figure 2.**
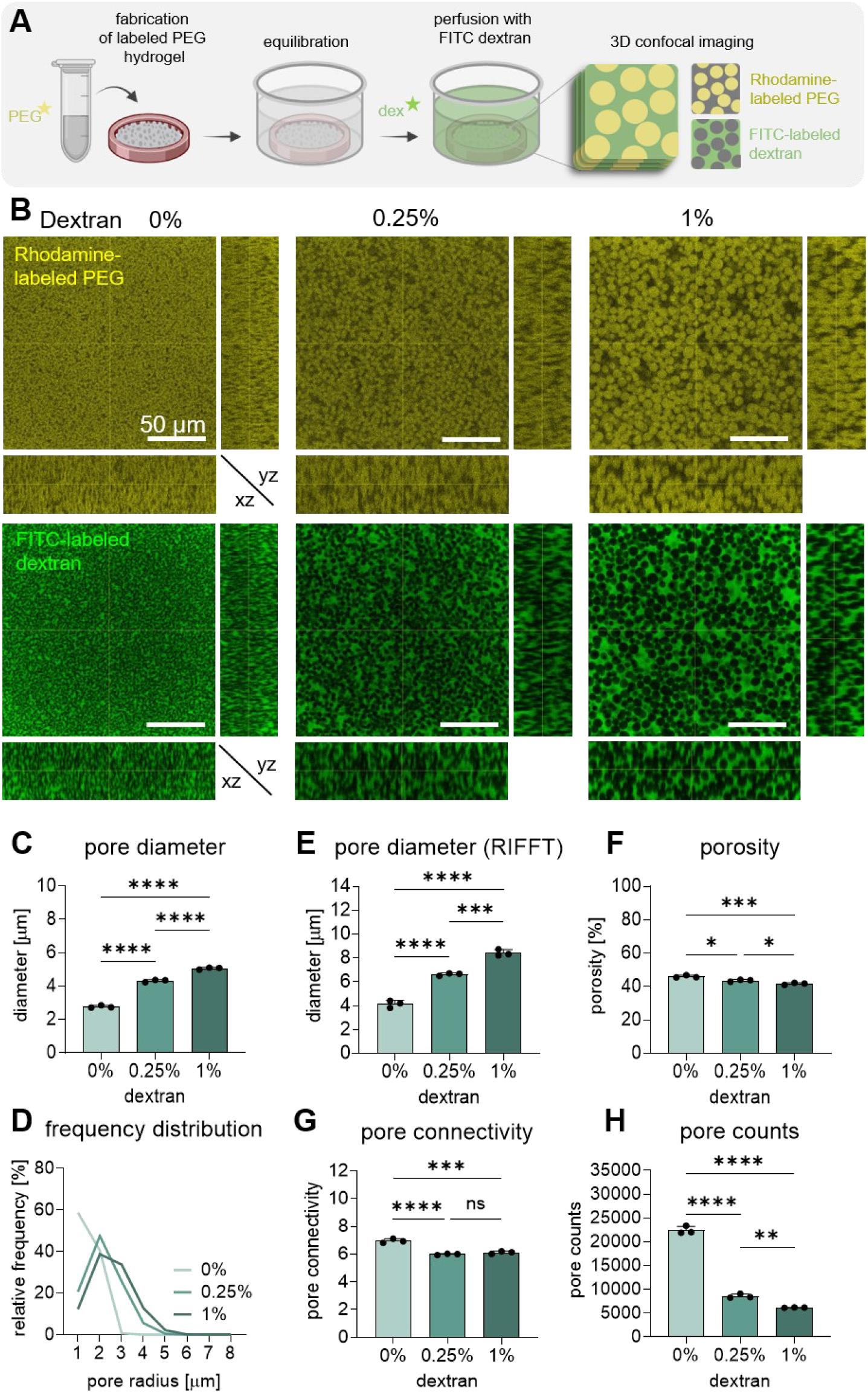
Characterization of acellular PEG hydrogels. **(A)** Schematic illustration of the fluorescence labeling process of acellular constructs using rhodamine-conjugated PEG and FITC-conjugated dextran to label void spaces. Illustration created with Biorender.com (https://biorender.com/djrsq27). **(B)** Orthogonal views of 50 μm Z-stack confocal images showing acellular rhodamine-conjugated PEG hydrogels fabricated with 0%, 0.25% or 1% dextran after perfusion with FITC-dextran. Scale bars, 50 μm. **(C)** Quantitative pore size characterization using 3D pore analysis ((**C**) pore diameter and frequency distribution, (**D**) porosity, (**E**) pore connectivity, (**F**) pore counts) and **(D)** FFT analysis (pore diameter - RIFFT) of acellular hydrogels using the FITC-channel. Each data point represents one hydrogel replicate with measurements averaged from 2 positions. In (C-G) each data point represents one hydrogel replicate with measurements averaged of 2 positions and 3 Z-planes. \**p* < 0.05, \*\**p* < 0.01, \*\*\**p* < 0.001, \*\*\*\**p*< 0.0001. Error bars represent the SD.

Next, we evaluated the physical properties of these hydrogels in terms of water-uptake and mechanical properties. The equilibrium water-uptake percentage decreases from ∼58% to ∼53% and ∼47% for hydrogels containing 0%, 0.25% and 1% dextran, respectively (**Supplementary Figure 1D**). Rheological characterization confirmed that all hydrogel formulations approached the storage moduli (G’) plateau following *in situ* crosslinking for 30 minutes at 37°C (**Figure 3A**). The G’ values at 30 min range from 180 Pa to 260 Pa (**Figure 3B**), confirming that all samples have a ultralow mechanical stiffness at the bulk level. The gelation time of three hydrogel formulations decreased from ca. 2.5 minutes to 1.5 minutes with increasing dextran concentration (**Figure 3C**). This acceleration may be attributed to the volume-excluding effect of dextran which may increase the local polymer concentration and lead to faster crosslinking.^26,27^ We note that beside pore size, gelation kinetics and water uptake are also influenced by the change of dextran concentration, reflecting the coupled thermodynamic and kinetic effects of dextran within this PIPS system. We reason that cells navigate the micropore space directly rather than the hydrogel network despite their biodegradability to cell-secreted MMPs. Our previous work revealed viscoelastic behavior in the described PIPS hydrogels due to the presence of HA.^13^ Since fast stress-relaxing alginate-collagen hydrogels have been shown to facilitate osteocyte morphogenesis,^20^ this viscoelastic nature of our HA-containing PEG microporous hydrogel may further enhance cellular morphogenesis. Together, these results demonstrate that increasing dextran concentration can modulate the kinetics of PIPS and also the pore size of PIPS hydrogels while keeping the mechanical stiffness in a comparably ultrasoft range.

**Figure 3.**
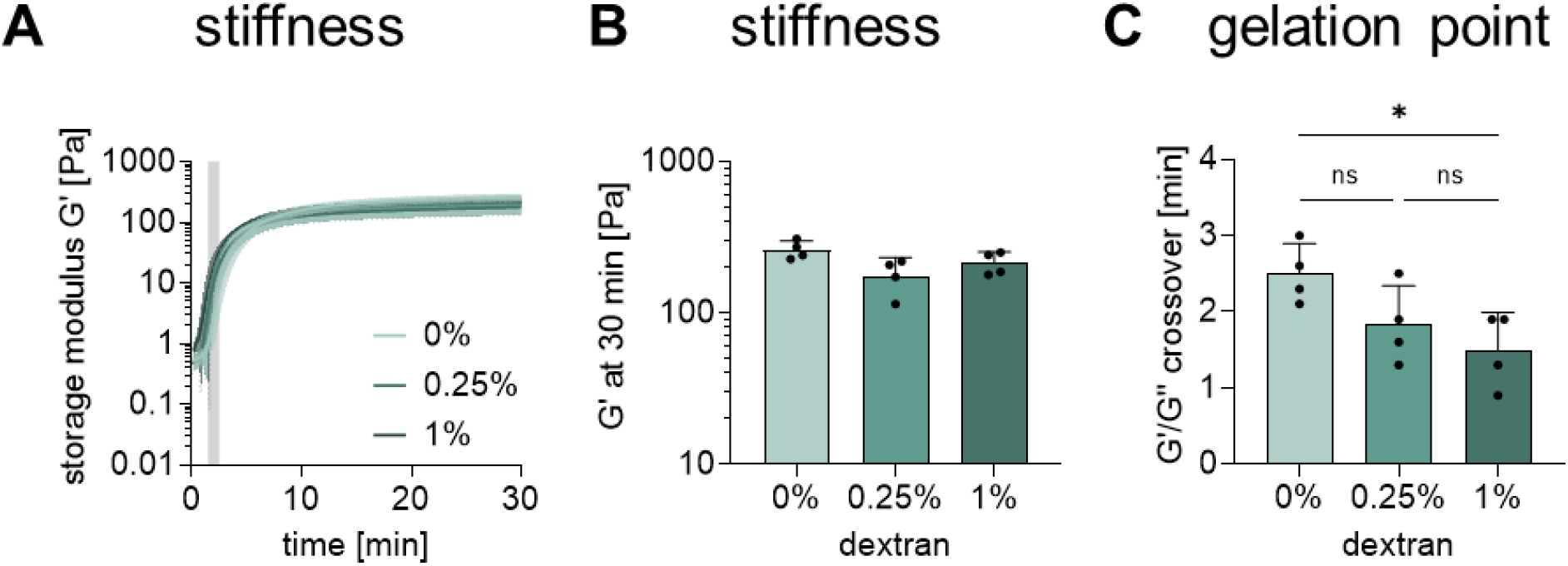
Rheological characterization of acellular PEG hydrogels. **(A)** Storage modulus G’ over time in a time sweep up to 30 min and **(B)** storage modulus plateau at 30 min of 0%, 0.25%, and 1% dextran hydrogels. **(C)** Gelation point (crosspoint G’>G’) of all three conditions. In A, each curve represents the average of 4 measurements, in B and C, each data point represents one measurement. *p* < 0.05. Error bars represent the SD.

### 2.2. Impact of HA digestion on microporosity and gel integrity

The presence of HA as a viscosity enhancer is essential for successful PIPS and formation of stable microporous hydrogels (**Figure 1B**).^13,23^ However, because HA may reside in the pore phase after gel formation, we evaluated whether its subsequent removal will alter the pore space. We compared passive diffusion during PBS equilibration with enzymatic treatment using hyaluronidase (HAase) into the hydrogels with varying amounts of dextran. The hydrogels were stained with both rhodamine-labeled PEG and FITC-labelled HA (**Figure 4A**) to visualize the hydrogel matrix and HA distribution, respectively. The presence of FITC-HA in the hydrogels was assessed at three time-points: directly after crosslinking, after initiation of the HA clearance protocols (PBS wash or HAase treatment) and after 20 hours of equilibration (**Figure 4B, Supplementary Figure 3**). No significant differences in HA fluorescence were found between the untreated and HAse-treated hydrogels at the initial timepoint. After 20h equilibration, the HAase-treated hydrogels exhibited almost complete HA removal in all hydrogel formulations, whereas very low levels of FITC-HA remained in untreated hydrogels (**Figure 4C**). The efficiency of HA clearance was highest in the hydrogels containing 1% dextran, suggesting that larger pores facilitate more efficient diffusive transport of HA from the hydrogel network.

**Figure 4.**
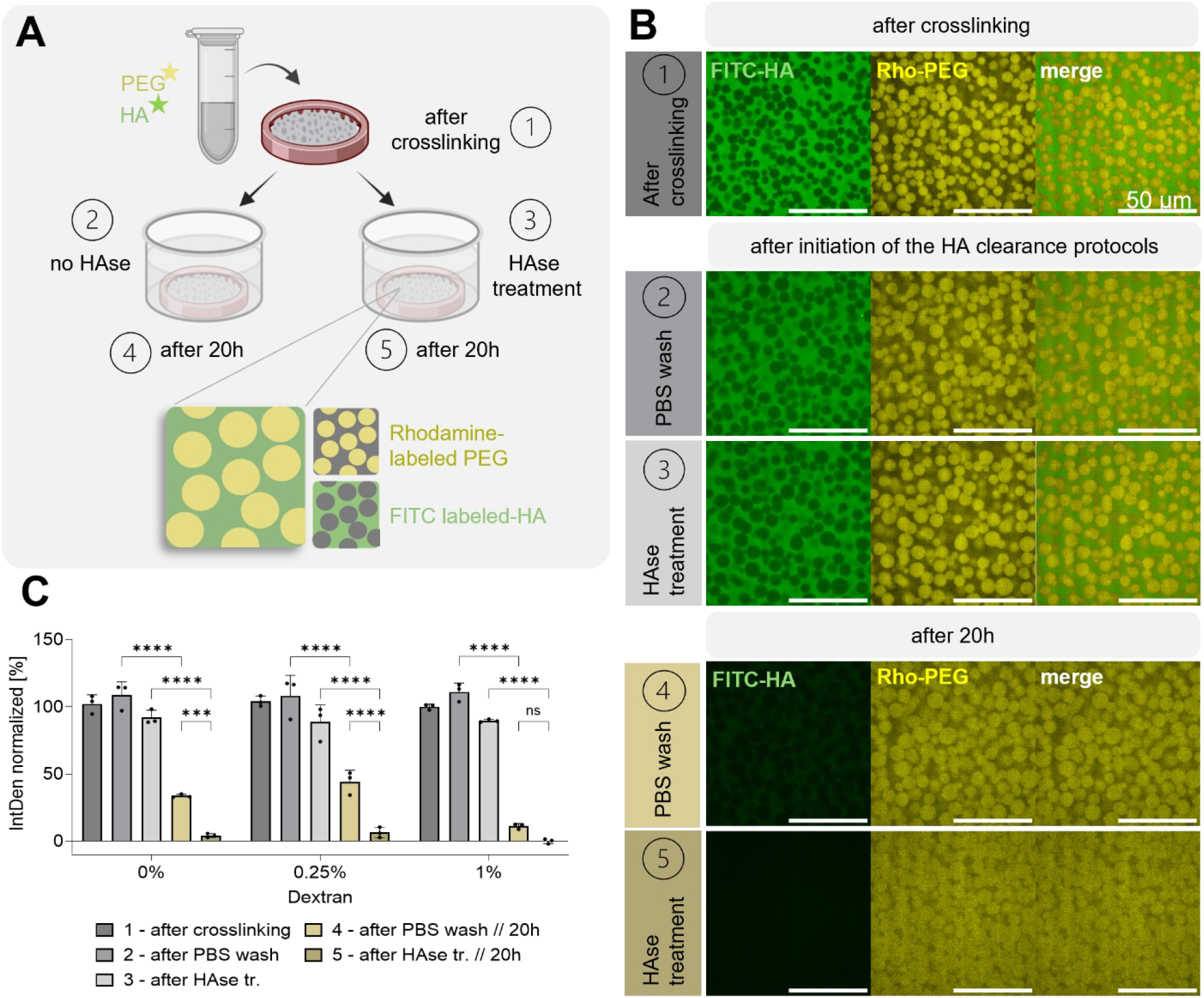
Comparison of passive and enzymatic HA clearance in microporous hydrogels. **(A)** Schematic illustration of the workflow comparing the standard wash protocol with PBS (2, 4) or enzymatic HAse digestion (3, 5) after crosslinking (1) of PEG hydrogels fabricated with Rhodamine labeled PEG and FITC labeled HA. Illustration created with Biorender.com (https://biorender.com/djrsq27). **(B)** Confocal images of (A) 1-5 of 1% dextran hydrogels. The intensity of the Rhodamine channel was automatically adjusted for better visualization of the hydrogel. Intensities in the FITC channel are the same for all conditions. Scale bars, 50 μm. **(C)** Quantification of signal intensities of FITC-HA confocal images (A) 1-5 of 1%, 0.25% and 0% dextran hydrogels. Values are normalized to 0% = (5) of 1% and 100% = (1) of 1%. Each data point represents one hydrogel replicate. Shown is the statistical analysis between (2) and (4), (3) and (5), and (4) and (5). \*\*\**p* < 0.001, \*\*\*\**p* < 0.0001, *ns* - non significant. Error bars represent the SD.

Although HAase treatment achieved near-complete HA clearance, it also altered pore size over time, indicating compromised hydrogel stability (**Figure 4B**). In contrast, the PBS wash was sufficient to remove most of the HA while preserving the pore sizes after 20 hours. These results demonstrate that PBS wash is sufficient to keep the microporosity accessible for ll culture.

### 2.3. Pore size influences cell viability and morphology during 3D osteogenic culture

We next investigated how fine-tuned pore size in PIPS hydrogels impacts on hOB cell viability and morphogenesis during 3D osteogenic cultures. hOBs were encapsulated at a constant density of 3×10^6^ cells/ml (**Figure 5A**) and cultivated under osteogenic differentiation conditions for up to 7 days. Live/dead assay after three days of cultivation confirmed high cell viabilities (>84%) for all conditions (**Figure 5B**). Notably, cell viability was slightly higher in hydrogels containing 0.25% and 1% dextran compared to the 0% dextran group, which is likely due to the improved nutrient transport throughout the porous space. This is consistent with another report on PVA-based microporous hydrogel via photo-PIPS.^14^ Morphological analysis using actin-nuclei staining revealed distinct cellular responses depending on the pore space. After 7 days of osteogenic culture, cells cultured in hydrogels containing 1% dextran exhibited enhanced cell spreading compared to those with smaller pores (0% dextran). Quantitative analysis of cell morphology revealed a 2-fold increase of dendrite length with increasing dextran concentration (**Figure 5C**). Total dendrite length measurements showed 3.5-fold higher values in hydrogels containing 1% dextran compared to those containing 0% dextran (Supplementary Figure 4). This suggests that larger pore dimensions provide greater spatial freedom for cytoskeletal organization and denrite formation. Together, these findings indicate that hydrogels with increased pore size enhances cell viability and promotes extensive 3D cellular morphogenesis. It is important to note that these microporous hydrogels are MMP-degradable, and thus cells may dynamically remodel the surrounding environments via proteolysis. ^13^

**Figure 5.**
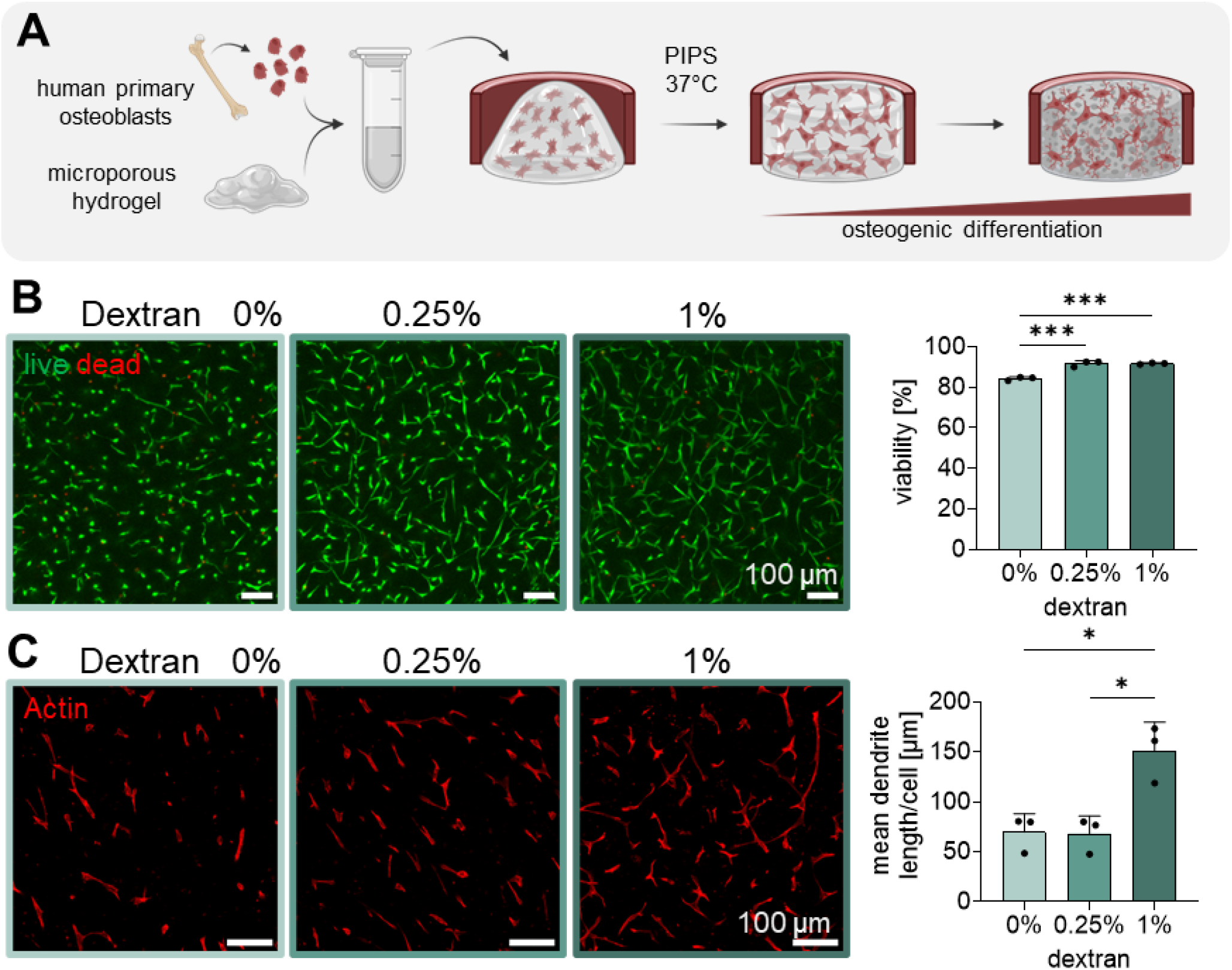
Analysis of cell viability and morphology in microporous PEG hydrogels. **(A)** Schematic illustration of the cell embedding process of patient derived primary osteoblasts in porous PEG hydrogels. Illustration created with Biorender.com (https://biorender.com/djrsq27). **(B)** MIP of 100 µm Z-stacks and quantification of live/dead viability assay of patient-derived osteoblasts embedded in 0%, 0.25% and 1% dextran PEG hydrogels after three days in osteogenic differentiation media. Each data point represents the average of 2 positions on one replicate. Error bars represent the SD. **(C)** MIP of actin staining (70 µm Z-stack) and quantification of the cellular morphology using NeuriteQuant^41^. Each data point represents one hydrogel replicate with measurements averaged of 2 positions. \*\**p* < 0.01, \*\*\**p* < 0.001, \*\*\*\**p* < 0.0001. Error bars represent the SD.

Consequently, the dynamics of cell-matrix remodeling within MMP-sensitive microporous PIPS hydrogels over time warrants further investigation.

### 2.4. 3D osteogenic differentiation and mechanosensing

To test whether pore size influences osteogenic differentiation, hOBs were encapsulated in hydrogels containing 0%, 0.25%, or 1% dextran and cultured under osteogenic conditions for up to 21 days. Osteogenic differentiation was quantitatively assessed by qPCR analysis of stage-specific marker genes (**Figure 6A**). The osteogenic marker alkaline phosphatase (gene: *ALPL*)^28^ exhibited a progressive upregulation over time across all conditions. Cells in the hydrogels containing 0% dextran showed the highest level of *ALPL* on 21 days, suggesting that spatial confinement may contribute to osteogenic differentiation. The mature osteoblast marker osteocalcin (gene: *BGLAP*)^29,30^ showed an initial upregulation within the first week followed by a moderate increase until day 21 across all groups. The expression of the early osteocyte marker Podoplanin (gene: *PDPN*)^31,32^ showed a similar increase over time. Cells in the hydrogels containing 0% dextran exhibited significantly higher levels of *PDPN* expression compared to the other two groups on day 21, implying that smaller pore geometries may accelerate the osteoblast-to-osteocyte transition. Collagen Type 1 (gene: *COL1A1*) expression increased initially and subsequently decreased by day 21, indicating the reduction in collagen synthesis as osteoblasts transition toward an osteocytic phenotype.^33^

**Figure 6.**
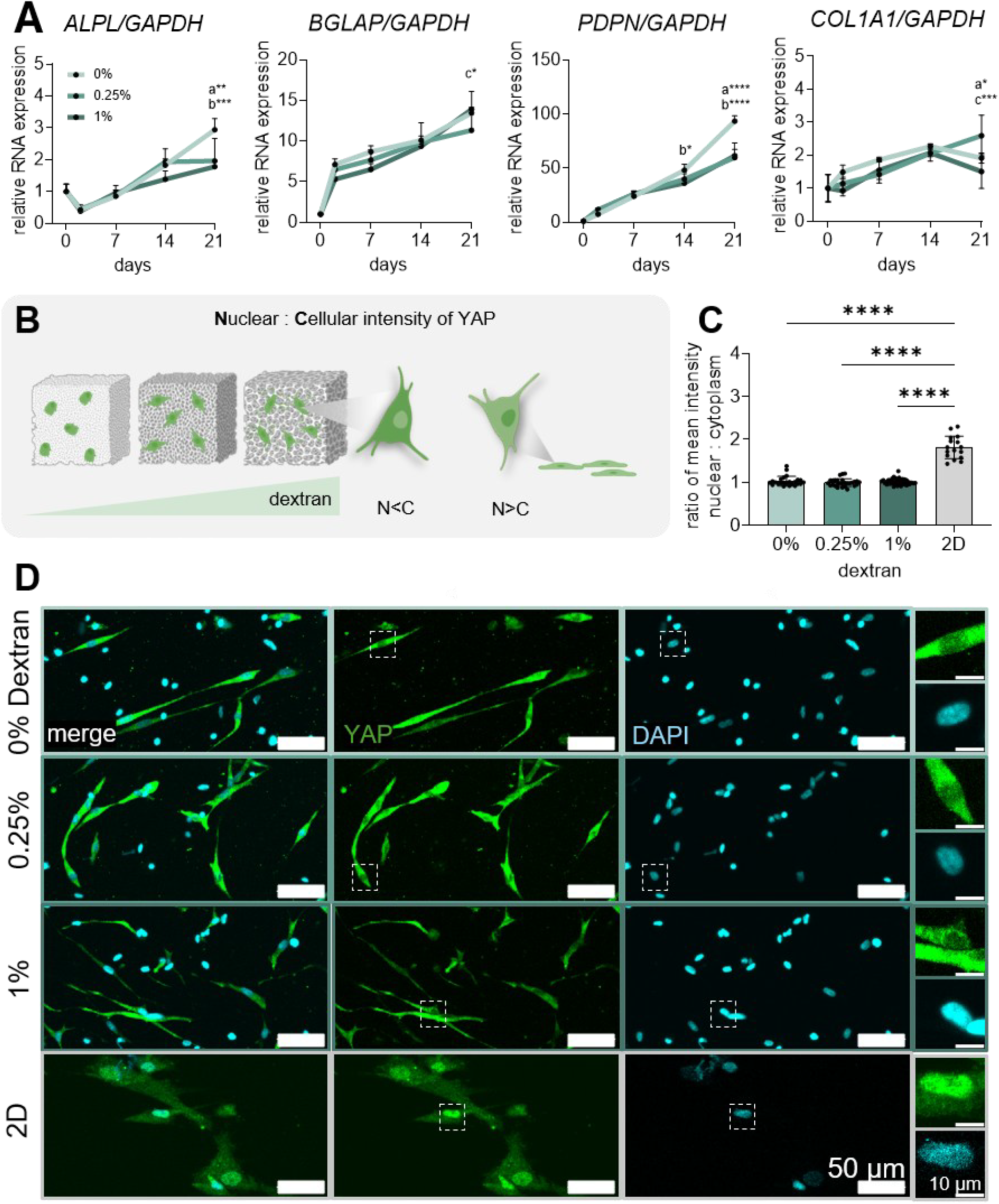
YAP localization and osteogenic gene expression of hOBs. **(A)** qPCR analysis of osteogenic marker during 21 days of hOB differentiation in 0-1% dextran PEG hydrogels. **(B)** Illustration of YAP expression in porous 0-1% dextran PEG hydrogels and **(C)** the quantification of the ratio of nuclear to cellular IntDen of YAP signal. Illustration created with Biorender.com (https://biorender.com/djrsq27). **(D)** Confocal images of immunofluorescence YAP staining and nuclear DAPI staining. Zoom-in regions are indicated by dashed lines. Scale bars, 50 µm and 10 µm. In (B) each data point represents 1 cell; data collected from 3 hydrogels per condition. In (A) each data point represents 1 hydrogel sample from one experiment. a) 0% vs. 0.25%, b) 0% vs. 1%, c) 0.25% vs. 1%. \**p* < 0.05, \*\**p* < 0.01, \*\*\**p* < 0.001, \*\*\*\**p* < 0.0001. Error bars represent the SD.

To investigate whether the observed pore size-dependent differences in osteogenic differentiation were associated with altered cellular mechanosensing, we next examined the subcellular localization of Yes-associated protein (YAP), a key mechanotransducer. After three days of culture, immunostaining of Yes-associated protein (YAP) revealed predominantly cytoplasmic localization across all groups. The nuclear:cytoplasmic (N:C) mean intensity ratio in all hOBs within 3D environments is significantly lower compared to those cultivated in 2D controls on glass (**Figure 6B-D**, Supplementary Figure 5). While YAP nuclear translocation is well established as a mechanotransducer in 2D,^34,35^ its behavior in 3D matrices, especially in biodegradable microporous hydrogels, is less well understood. Increasing evidence suggests that YAP activation is influenced by multiple factors beyond stiffness alone, including matrix degradability, cell dimensionality and cell spreading.^35^ Several recent studies have suggested that soft 3D matrices favor cytoplasmic YAP retention.^36–38^ Notably, within a given stiffness range, pore geometry has been shown not to significantly affect YAP activation^38^. Our findings indicate that the selected microporosities in ultrasoft PIPS hydrogels do not activate YAP. Future work is required to identify alternative pathways that may clarify the effect of fine-tuned microporosities on osteogenic progression.

Osteogenic differentiation of MSCs has been widely demonstrated on stiff hydrogel substrates (10– 50 kPa)^3,39^, yet these studies primarily address 2D cultures and uncommitted progenitor cells. In contrast, our results emphasize that 3D embedded primary hOB cells undergo osteogenic differentiation within soft microporous hydrogel environments where pore size and biodegradability of the matrix can be modulated, both of which may contribute to osteogenic progression.^13^ Müller et al.^14^ showed the importance of pore space for osteogenic differentiation in RGD-functionalized, PVA-based microporous hydrogels that are only hydrolytically degradable. Zauchner et al. recently demonstrated that MMP-sensitive microporous PEG hydrogels support 3D bone cell network formation and osteogenic differentiation.^13^ While most other reported matrices are either stiffer, nanoporous or feature macroporosity with pore sizes orders of magnitudes larger,^40–46^ direct comparison of pore geometry effects on osteogenic differentiation remains difficult. Mechanistically, reduced nutrient diffusion within hydrogels with smaller pores may generate a locally hypoxic microenvironment that may promote osteogenic lineage commitment,^1,40,47^ potentially reinforcing the effect of increased cell–wall contact as dendritic processes extend into the confined pore space. Although these effects became evident at day 21, further investigation with a longer culture duration is warranted. Taken together, these results imply that even a minor change of pore size in the selected microporous PIPS hydrogel modulates osteogenic differentiation of primary hOBs.

### 2.5. Cell infiltration into microporous hydrogels

For potential *in vivo* applications, we finally investigated whether these microporous hydrogels can support cell infiltration. Human mesenchymal stem cells (hMSCs), owing to their migratory capacity, ^48^ were top-seeded onto the surface of equilibrated hydrogels (**Figure 7A**) and their infiltration was monitored by time-lapsed live-cell tracking. To better understand the mechanism of cell infiltration, we included an RGD-free control group for the hydrogels containing 1% dextran. After four days of osteogenic differentiation, cells atop the 0% dextran hydrogels showed negligible horizontal infiltration. Cells in the hydrogels containing 0.25% dextran showed moderate invasion where the tip cells cleared a path while keeping connected to the cells behind. In contrast, cells in hydrogels containing 1% dextran extensively infiltrated the porous space, demonstrating that microporosity facilitates cell migration in a pore size-dependent manner (**Figure 7B**). Notably, cells in RGD-free hydrogels showed no signs of infiltration. This contrasts with neural cells, where microporous PEG hydrogels support neurite extension without adhesion cues, ^23^ and is consistent with the necessity of RGD motifs for HDF spreading in PIPS hydrogels by Müller et al.^14^ Together, these data suggest that cell infiltration into synthetic microporous hydrogels requires both cell adhesion and cell-scale microporosity, which may find future *in vivo* bone regeneration applications.

**Figure 7.**
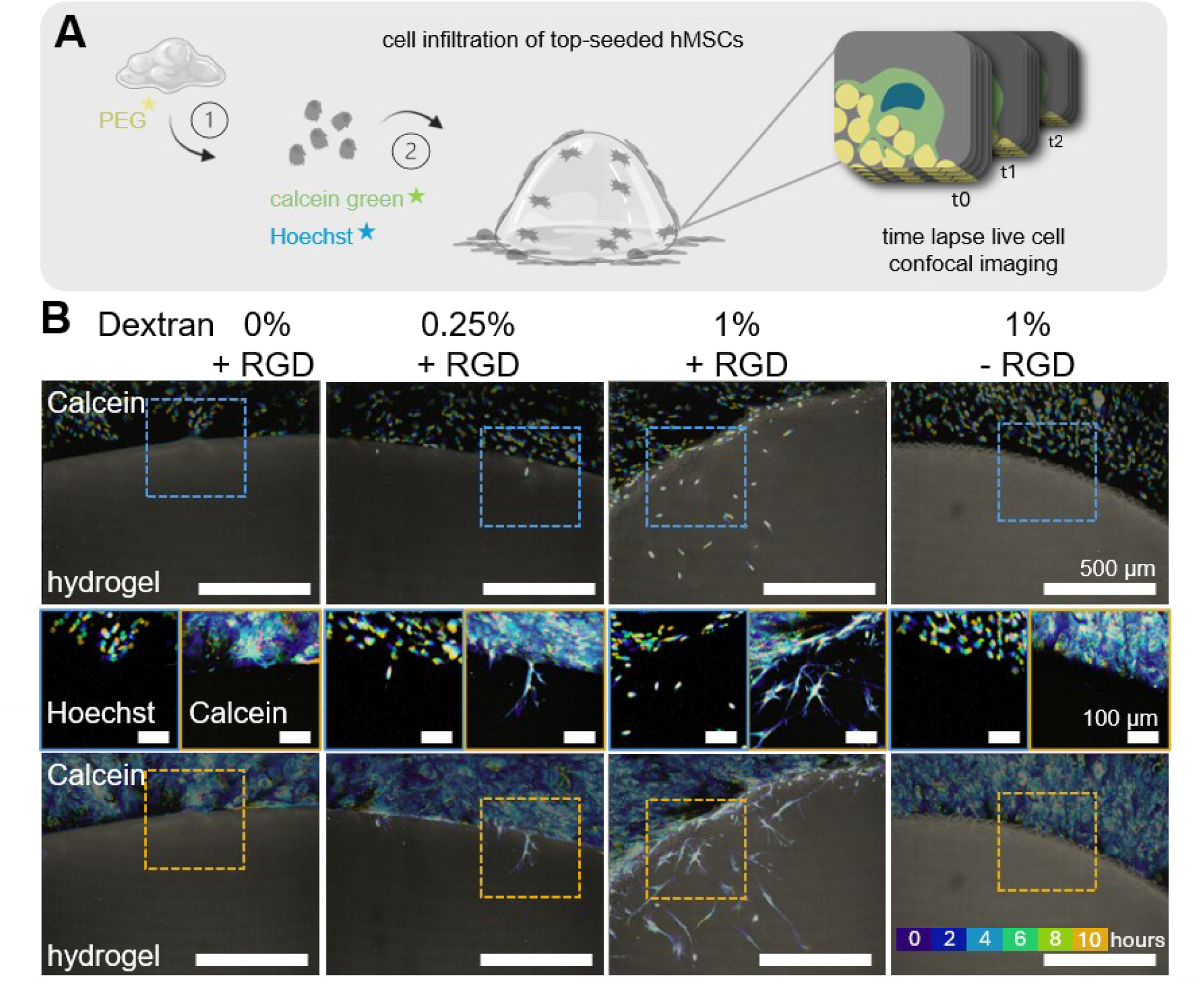
Cell infiltration of top-seeded hMSCs into microporous hydrogels. **(A)** Schematic illustration of the analysis of cell infiltration of top-seeded cells. Illustration created with Biorender.com (https://biorender.com/djrsq27). **(B)** Temporal color-coded analysis of nuclei (Hoechst), cell bodies (calcein) and hydrogel (rhodamine-labeled 4-PEG-VS) of top-seeded hMSCs after 4 days culture on different hydrogels. Scale bars, 500 μm (up, bottom) and 100 μm (middle).

## 3. Conclusions

In conclusion, we investigated how fine-tuned cell-scale microporosities influence bone tissue morphogenesis and differentiation *in vitro* using a modular synthetic microporous hydrogel. By tuning dextran concentration, we generated hydrogels with varying cell-scale pore sizes while maintaining comparable mechanics, adhesiveness and biodegradability. Even with a minor change of dextran concentration, we observed pore size-dependent effects: hydrogels with larger pores promote cell viability, dendritic morphogenesis, and hMSC cell infiltration, whereas hydrogels with smaller pores enhance osteogenic commitment, possibly due to spatial confinement. This current PIPS system creates pore volumes up to ∼65 μm^3^ (∼5 μm in diameter) through fine-tuned phase separation while maintaining ultralow stiffness (<300 Pa), enabling investigation of cell-material mechanobiology within an osteoid-like soft environment. This platform uniquely allows the study of the effects of 3D spatial confinement on osteogenic differentiation. Future studies could explore substantially larger pore sizes (>20 μm). However, at these dimensions, bone cells likely to spread along the pore surface rather than fully encapsulated, resulting in cell–matrix interactions and curvature sensing that are governed by fundamentally different mechanobiological principles than those operating in true 3D embedding. Nevertheless, uncovering the mechanistic differences in cell-matrix mechanosensing and tissue differentiation between healthy and patient-derived bone cells may pave the way for personalized drug screening and novel therapeutic strategies.

## Author contributions

Conceptualization: X.-H.Q., M.H.; Methodology: M.H., D.Z.; Formal analysis: M.H., D.Z.; Resources: X.-H.Q., R.M., C.G., M.R.; Data curation: M.H., D.Z., E.K.; Writing original draft: M.H.; Review and editing: X.-H.Q., M.H., D.Z., M.E., R.M., P.J.L., C.G., M.R.; Visualization: M.H.; Supervision: X.-H.Q., R.M.; Project administration and funding acquisition: X.-H.Q.

## Acknowledgements

This work was supported by the Swiss National Science Foundation (SNSF) through the National Research Program “Advancing 3R – Animals, Research and Society’ (NRP 79, grant no. 206501), SNSF project funding (grant no. 207542), and ETH grant (grant no. 24-1 ETH-060). The authors thank the Scientific Center for Optical and Electron Microscopy (ScopeM) of ETH Zurich for providing the microscopy facilities. Further, the authors thank Esther Rosenwald for her assistance in the culture maintenance.

## TOC

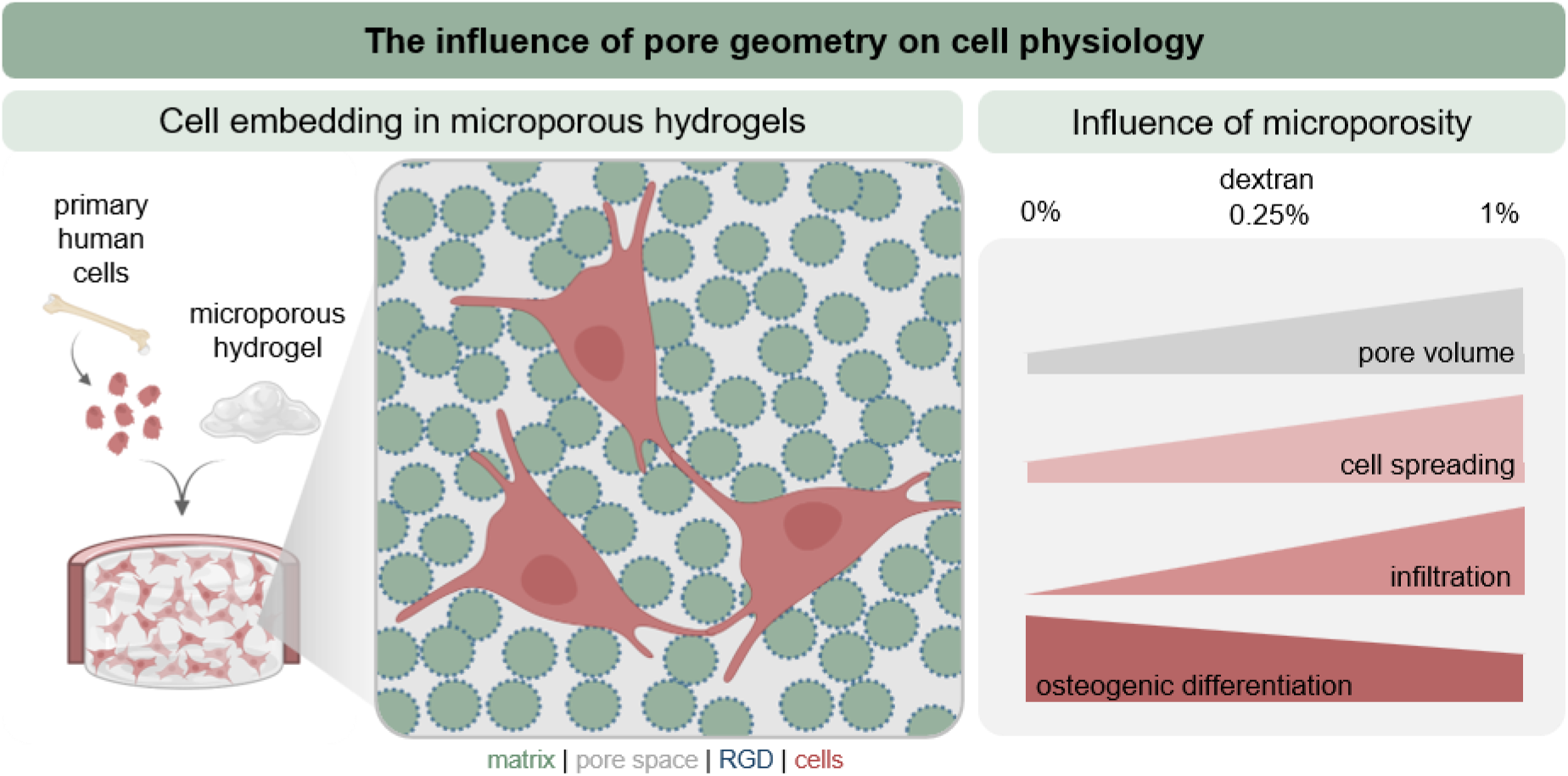

