## Supplementary Information for "A Modular Synthetic Hydrogel with Cell-scale Micropores Modulates Osteocyte-like Morphogenesis and Osteogenic Differentiation *In Vitro*"

List of Content
- Experimental section
- Supplementary Table 1-2
- Supplementary Figure 1-5
- References

### Experimental section

#### Synthesis of 4-arm-polyethylene glycol (PEG) and rhodamine labeled PEG

Rhodamine-labeled and unlabeled 4 arm-PEG-vinyl sulfone (4-PEG-VS) were synthesized as described by Broguiere et al., 2019 [1], and Zauchner et al., 2024 [2]. For both PEG variants, 4-PEG-thiol was reacted with divinyl sulfone in TEOA buffer. For Rhodamine-labeled 4-PEG-VS, tetramethyl rhodamine-5-maleimide was pre-conjugated to 4-PEG-thiol before reaction with divinyl sulfone. After sterile filtration, aliquoting and lyophilization, both PEG variants were stored at –20 °C. Shortly before usage, PEG was dissolved through vortexing and sonication in sterile filtered 0.3M HEPES buffer pH 7.4 and stored at 4 °C.

#### Characterization of pore size, porosity and connectivity

PEG hydrogels were cast as described in *Synthesis of 4-arm-polyethylene glycol*. After swelling for 1 day in PBS, hydrogels were incubated for 4 days in 2 mg/ml FITC-dextran 2 MDa in PBS while shaking at room temperature for 100 rpm. Thereafter, hydrogels were imaged with a Leica SP8 confocal microscope. For 3D pore quantification, Z-stacks of 50 µm were taken. The confocal images were analyzed using the MATLAB algorithm by Vandaele et al.[3], which segments the images and approximates pores as spheres to quantify beside other parameters pore size, pore connectivity, and porosity. An adaptive threshold sensitivity of 0.6 was applied. For each of three independent samples, two Z-stacks on two different positions were acquired and analyzed and subsequently averaged.

#### Rheology

Rheological measurements were performed as described before with some adjustments.[2] Briefly, hydrogels were prepared on ice. 70 µl of gel precursor was then loaded onto a glass-bottomed Anton Paar MCR302 rheometer (82868246) with a PP20 plate, covered with mineral oil and the storage modulus (G') was monitored at 37°C over 30 min in a time-sweep experiment (gap: 100 µm, frequency: 1 Hz, strain: 5%).

#### Effect of HAse treatment

Hydrogels were prepared as described before, with the exception of using FITC-HA (1:50) and Rhodamine labeled PEG (1:20). One set of hydrogels was imaged directly after crosslinking, the second set was washed with PBS, imaged and incubated in PBS overnight and the third was washed, incubated 1h at 37 °C in 1 mg/ml hyaluronidase (HAse) in PBS, washed with PBS, imaged and incubated in PBS overnight. Hydrogels were imaged again the following day (after 20h). Imaging was performed using an Avantor BC43 spinning disk confocal microscope at 63x magnification. Rhodamine PEG channel is shown as auto-adjusted intensity for better visualization. Fluorescence intensities of the HA channel were standardized and quantified using the integrated density (IntDen) measurement in Fiji2.

### Osteogenic differentiation of primary human osteoblasts

After cell harvesting, cells were counted for a final concentration of $3\times10^6$ cells/ml and resuspended in HA. All other components were combined on ice and mixed. HA-cell mix was added to the PEG mix and vortexed again. As last step, the crosslinker was dissolved, filtered and added to the mix. After vortexing, 15-20 µl of hydrogel mix was immediately cast into previously sterilized 4 mm in diameter silicon molds on cover glasses cooled on ice. The hydrogels were then crosslinked in the incubator at 37 °C and 5% $CO_2$ for a minimum of 45 min and washed with PBS. Embedded hOBs were differentiated using osteogenic differentiation media (OGM) composing of Low-Glucose DMEM (phenol red free), 10% Fetal bovine serum (FBS), 1% antibiotics/antimycotics (AntiAnti), 25 mM HEPES and freshly added 10 mM b-glycerolphosphate (bGP, Acros, 410991000) and 50 µg/ml ascorbic acid and kept at 37 °C and 5% $CO_2$. Cultures were kept up to three weeks while media was changed 3 times per week. On day 2, 7, 14, and 21 hydrogels for RNA isolation were washed with PBS and collected using a positive displacement pipette, shock frozen in liquid $N_2$ and stored on –80 °C until further processing. Samples for immunohistochemistry were washed with PBS, fixed in 4% paraformaldehyde (PFA) (Sigma-Aldrich, 15-812-7) for 45 min (3D samples) or 15min (2D samples), washed again with PBS and stored at 4°C until further processing.

### Cell infiltration

Sterile rhodamine-labeled PEG hydrogels with 0%, 0.25%, 1% and 1% without RGD dextran hydrogels were prepared. The gels were equilibrated for 1 day in hMSC differentiation media. hMSCs were harvested at P6 and ~ 30,000 cells/cm$^2$ were stained with NucBlue Live ReadyProbes Reagent (Thermo Fisher, R37605) and 1:500 Calcein Green AM (Sigma-Aldrich, 56496) in PBS for 30 min. Cells were top seeded and incubated for 4 hours before imaging on the BC43 Benchtop Confocal Microscope with an incubator chamber. After 5 days, cells were re-stained with NucBlue and Calcein Green AM and imaged over 10h, with every second hour a Z-stack. LUT adjusted maximum intensity projection (MIP) of Z-stacks were used for temporal color-coded analysis of the NucBlue and Calcein Green AM channel in Fiji2. Samples were fixed and stored at 4°C until further use.

### Hydrogel composition and acellular hydrogel fabrication

For details on the reagents, see **Supplementary Table 1**. Dextran and RGD peptide were dissolved and sterile filtered using a 0.22 µm filter. HA was sterilized using UV treatment in the laminar flow and PEG precursor was prepared sterile as described in *4-arm-PEG synthesis*. After combining all components except crosslinker, the PEG mix was cooled on ice. The crosslinker was prepared just on time, sterile filtered and added to the PEG mix. 15-20 µl hydrogel was cast into a cylindrical silicon mold (mold diameter: 4 mm, mold height: 2 mm) in a confocal dish which was placed on an ice-cooled metal plate for even cooling before crosslinking. The samples were transferred into the incubator (37 °C, 5% $CO_2$) on a preheated metal plate for even heat distribution. After a minimal 45 min of crosslinking, samples were washed with PBS and let swell for at least 24 h in PBS before further analysis.

### Characterization of pore size using 2D Fast Fourier transform

As alternative pore quantification method, we used the 2D Fast Fourier Transform (2D-FFT)[4]. Raw images were preprocessed using a bandpass filter before analysis. Images were then converted to grayscale and processed using a Hamming window before applying the 2D-FFT. Radial intensity of the FFT (RIFFT) were extracted from the frequency data through radial averaging, generating intensity curves. Peaks were manually determined from these curves, and corresponding length scales were calculated as $q^{-1}$. For each of three independent samples, images were acquired at three positions and across three different z-planes, and subsequently averaged.

### Swelling assay

Hydrogels were cast as described in *Hydrogel fabrication.* Hydrogels were weighted directly after crosslinking (Hd) on a fine balance and again after 24h swelling in PBS at 37 °C (Hw). The swelling ratio (SR) was calculated as follows: SR = (Hw-Hd)/Hd

### Primary human osteoblast expansion

Primary human osteoblasts were processed and provided from Kinderspital Zürich. Once obtained a frozen vial, cells were cultivated in osteoblast expansion media (OEM) composing of High-Glucose DMEM (with sodium pyruvate and L-Glutamine), 10% Fetal bovine serum (FBS, Gibco, Lot#42F7190K or 2440094), 1% antibiotics/antimycotics (AntiAnti, Gibco, 15240062), 10mM HEPES (Gibco, 15630080**)** and freshly added 100 μg/ml ascorbic acid (Sigma-Aldrich, A92902-100G) and kept at 37 °C and 5% $CO_2$. Media was changed 3 times per week. Primary osteoblasts were used up to Passage 6.

### Live/dead assay

Embedded cultures were washed 3 times with PBS, incubated with 1:500 Ethidium-homodimer-1 (EthD-1, Sigma-Aldrich, 460439) and 1:1000 Calcein Green AM (SigmaAldrich, 56436) for 1 h in PBS while shaking, washed with PBS and kept in culture media until imaging. Samples were immediately imaged on a Zeiss confocal Microscope LSM780 with 10x magnification. Z-Stacks of 100 µm were used for live and dead quantification using Image J.

### Fluorescence and immunofluorescence staining

For the immunofluorescence staining, the procedure was followed as outlined in Piccolo lab[5] protocols with some adjustments. Briefly, the fixed samples were permeabilized with 0.3% Triton X-100 (Sigma-Aldrich, 9002-93-1) in PBS for 20 minutes, followed by blocking in 10% goat serum in PBS-T (0.1% Triton X-100 in PBS) for 3 hours. After blocking, the samples were washed three times with PBS-T and incubated overnight at 4°C with the primary antibody (YAP) in 1% bovine serum albumin (BSA, Sigma-Aldrich, 9048-46-8), 2% goat serum and 0.1% Triton X-100 in PBS. The following day, the samples were washed three times with PBS-T and then incubated with the secondary antibody (1:500), phalloidin 647 (1:400) (Invitrogen, A30107), and DAPI (1:1000) (ThermoFisher, 62248) for 4 hours at room temperature. After an additional three washes with PBS-T, the samples were stored in PBS at 4°C until imaging at the Avantor BC43 Benchtop Confocal Microscope. For actin/nuclei staining only, the protocol was simplified to permeabilization and continuing directly with washing and staining.

### RNA isolation

Hydrogels were thawed on ice and homogenized using a Fisherbrand™ Pellet Pestle™. 300 µl of Trizol (Invitrogen, 15596018) was added to each sample and further homogenized. Another 700 µl of Trizol was added and incubated for some minutes. 300 µl Chloroform (Sigma-Aldrich, C2432-500ML) was added and vortexed. The solution was added to Phasemaker™ Tubes (Invitrogen, A33248) and centrifuged at full speed for 20 minutes. Afterwards, the clear phase was collected and mixed 1:1 with 70 % (v/v) Ethanol before transferring on a Qiagen RNeasy Micro Kit column (Qiagen, 74004). RNA was further isolated using the Qiagen RNeasy Micro Kit protocol, and RNA was eluted in 20 µL $H_2O$. After isolation, RNA was stored at -70°C.

### Reverse transcription and quantitative gene expression analysis (qPCR)

A total of 270 ng of RNA was reverse transcribed into cDNA using the iScript™ Reverse Transcription Supermix for RT-qPCR (Bio-Rad, 1708840) according to the manufacturer's protocol. For each qPCR reaction, 5.5 ng of the resulting cDNA was used as input. Reactions were prepared using the GoTaq® Probe qPCR Master Mix (Promega, A6101) and TaqMan™ Gene Expression Assay primers (see **Supplementary Table 2**). For relative quantification, a 1:2 serial dilution (v/v) (1, 1/2, 1/4, 1/8, 1/16) of a standard sample was included. All samples and standards were run in duplicates. The qPCR was performed on a CFX96™ Real-Time System (C1000 Touch™ Thermal Cycler) with initial denaturation at 95°C for 2 minutes, followed by at least 40 cycles of 95°C for 30 seconds and 60°C for 2 minutes. Gene expression levels were normalized to the housekeeping gene *GAPDH*.

### Quantification of cell morphology

To quantitatively assess cell morphology, hOBs stained for F-actin and nuclei were imaged using a Leica SP8 confocal microscope with a 20× objective. Z-stacks of 70 µm were obtained and maximum intensity projections (MIPs) were generated. Quantification of total dendrite length per image and mean dendrite length per cell (normalized to nuclei count) was performed using the ImageJ plugin NeuriteQuant [6]. Analysis parameters were optimized as follows: neurite detection width = 9, neurite detection threshold = 25, neurite cleanup threshold = 10, and neuronal cell body detection threshold = 7.

### YAP quantification

For each sample at 2 positions, a Z-stack of 200 µm was acquired on the Avantor BC43 Benchtop Confocal Microscope. Image analysis was performed in Fiji 2/ImageJ with ROIs of the cytoplasm (YAP channel – $ROI_{cytoplasm}$) and nucleus (DAPI channel - $ROI_{nucleus}$). The N:C ratio was calculated using the mean intensity of $ROI_{nucleus}$: $ROI_{cytoplasm}$.

### Statistical analysis

Statistical analysis was performed using GraphPad Prism version 10. Data were analyzed using one-way or two-way ANOVA followed by Tukey's multiple comparisons test, as appropriate. Dendrite length data were assumed to follow a lognormal distribution. A one-way ANOVA with Tukey's multiple comparisons with a single pooled variance was performed on log-transformed values (GraphPad Prism lognormal ANOVA option). For comparisons between three groups, an ordinary one-way ANOVA followed by Tukey's multiple comparisons test with a single pooled variance was applied. For the HA treatment experiment and gene expression analysis an ordinary two-way ANOVA followed by Tukey's multiple comparisons test with a single pooled variance was applied. Significance levels are indicated as follows: *$p < 0.05$, **$p < 0.01$, ***$p < 0.001$, ****$p < 0.0001$. Error bars represent the standard deviation (SD). For all cellular experiments, each data point represents one independently cultured sample from a single well within one experiment.

### Supplementary Table 1

| reagent | | stock for hydrogel preparation | final |
|---|---|---|---|
| 4-PEG-VS or Rhodamine-labeled 4-PEG-VS | 2.0 kDa | 20% in 0.3M HEPES buffer | 2% |
| PBS pH 7.4 | | | |
| Dextran (Sigma-Aldrich 31392-10G) | 500 kDa | 8% or 16% in PBS pH 7.4 | 0-4% |
| Hyaluronic acid (HA) (Sigma-Aldrich, 9067-32-7) or FITC-HA (TdB labs, Batch No. 21127) | 1.5 MDa | 1% in low-Glucose DMEM (phenol red) (Gibco, 41966-052) | 0.525% |
| Crosslinker (China Peptides) | N-C: GCRDGPQGIWGQDRCG | 6.23% in 3M TEOA pH 8 | 0.8 SH/ene |
| RGD peptide (China Peptides) | N-C: GRCGRGDSPG | 1.38% in PBS pH 6 | |

**Supplementary Table 1** Reagents and concentrations for PEG hydrogel preparation.

**Supplementary Table 2**

| gene | TaqMan™ Assay |
|---|---|
| Human *GAPDH* | Hs02786624_g1 |
| Human *ALPL* | Hs01029144_m1 |
| Human *PDPN* | Hs00366766_m1 |
| Human *COL1A1* | Hs00164004_m1 |
| Human *OCN* | Hs01587814_g1 |

**Supplementary Table 2** TaqMan™ Gene Expression Assay primer used for qPCR analysis of osteogenic differentiation.

**Supplementary Figure 1**

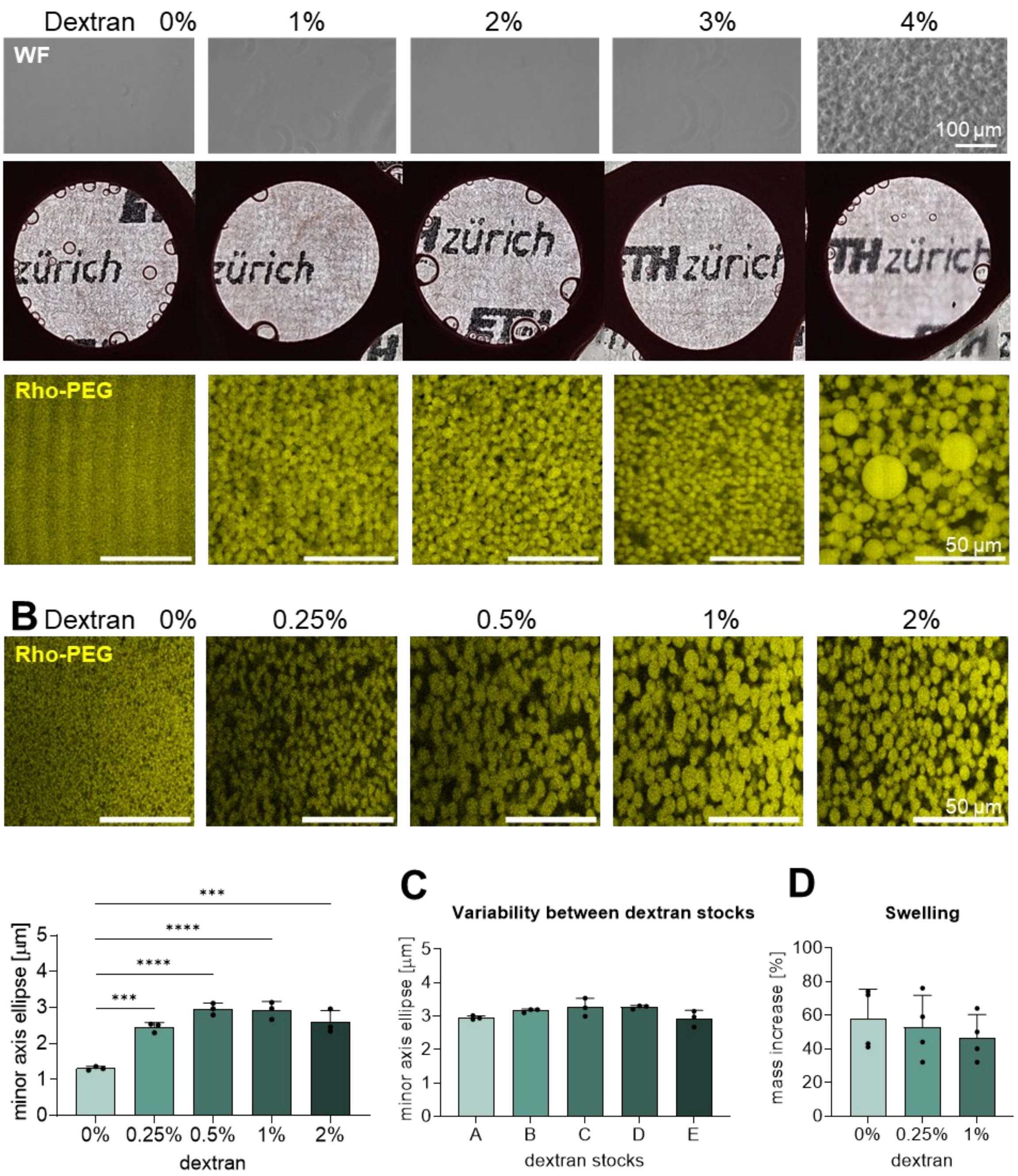


**Supplementary Figure 1 (A)** Wide field (top) and macroscopic (middle) images of PEG-dextran hydrogel compositions (0-4% dextran) without crosslinker. The ETH Zurich logo is used to observe turbidity of hydrogel compositions. Confocal images showing the pore structure of the same PEG-dextran formulations with crosslinker. **(B)** Confocal images of PEG-dextran formulations from 0-2% dextran and the quantification of pore size diameter using ImageJ ellipse fit analysis in Fiji2. **(C)** Variability between 1% dextran formulations using five different dextran stocks (A-E). Pore diameter is analyzed using ImageJ ellipse fit analysis. **(D)** Swelling ratio of 0-1% dextran hydrogels.

**Supplementary Figure 2**

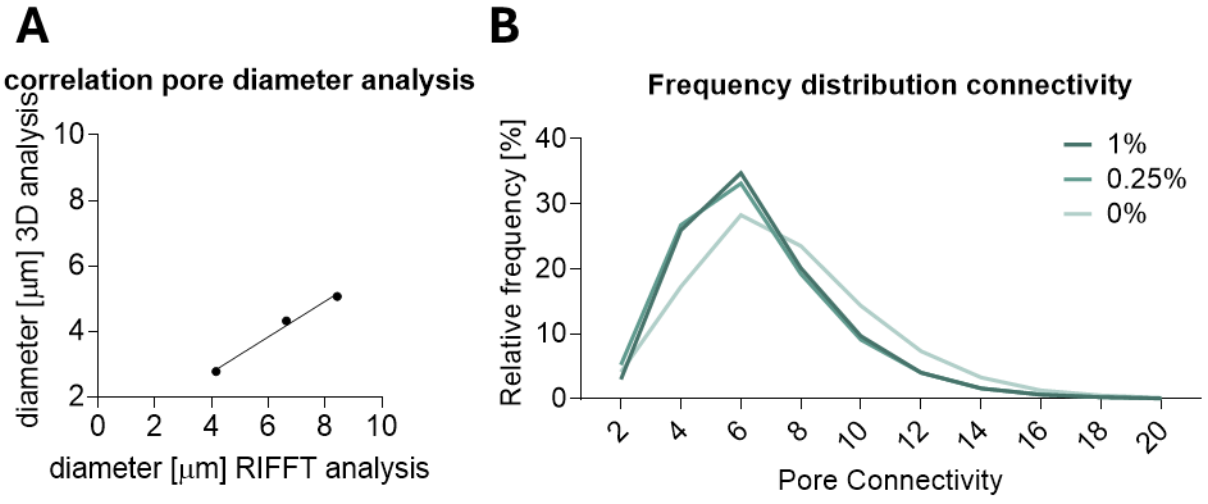


**Supplementary Figure 2 (A)** Correlation between 3D and RIFFT pore size analysis. **(B)** Frequency distribution of pore connectivity analyzed by 3D pore size analysis.

**Supplementary Figure 3**

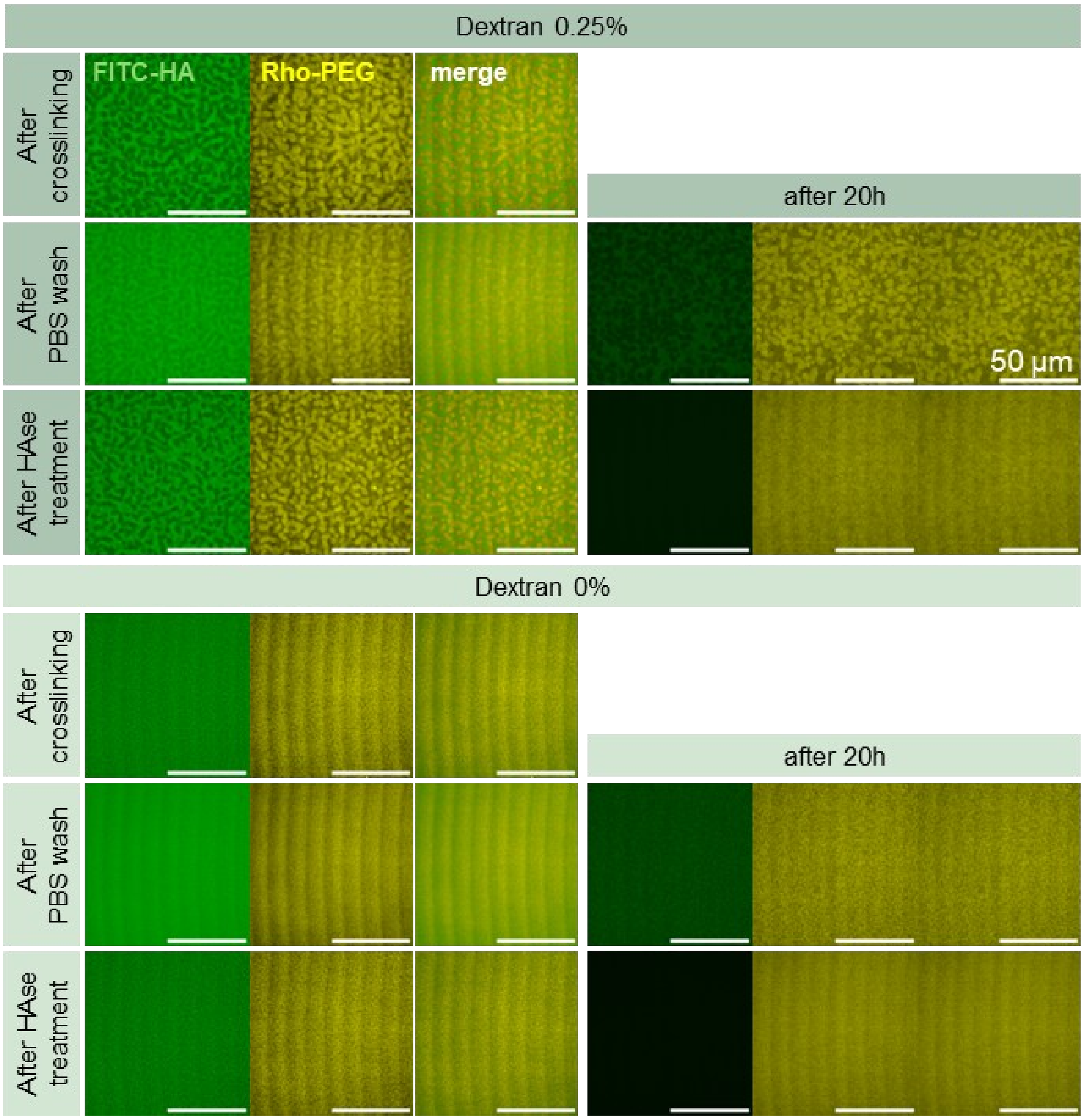


**Supplementary Figure 3** Confocal images of **Figure 3A** 1-5 of 0% and 0.25% dextran hydrogels. The intensity of the Rhodamine channel was automatically adjusted for better visualization of the hydrogel. Intensities in the FITC channel are the same for all conditions.

**Supplementary Figure 4**

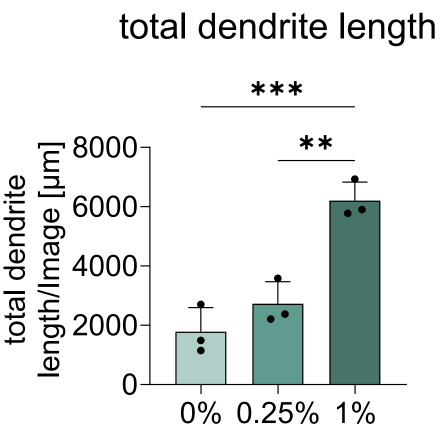


**Supplementary Figure 4** Quantification of cellular morphology using NeuriteQuant[6]. Each data point represents one hydrogel replicate with measurements averaged of 2 positions. Shown is the mean +SD.

### Supplementary Figure 5

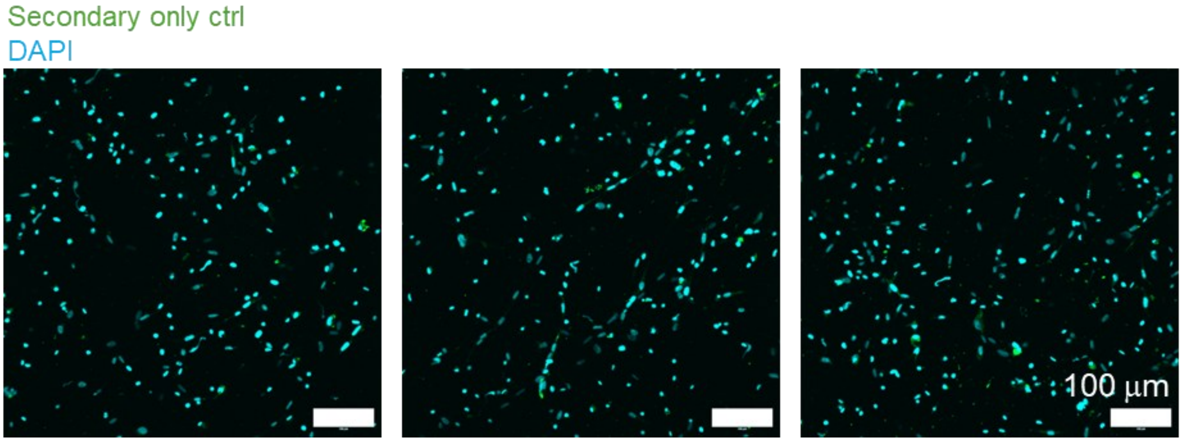


**Supplementary Figure 5** Confocal images of maximum intensity projections of 200 µm z-stacks of secondary only control of YAP staining.
